# Alterations of gut microbiota in Down syndrome and their association with Alzheimer’s disease

**DOI:** 10.64898/2026.04.03.716276

**Authors:** Camilla Pellegrini, Francesco Ravaioli, Sara De Fanti, Claudia Sala, Magali Rochat, Virginia Pollarini, Polischi Barbara, Alberto Pasti, Margherita Grasso, Maria Rambaldi, Francesco Cardoni, Nicola Grotteschi, Filippo Caraci, Pietro Cortelli, Federica Provini, Raffaele Lodi, Luca Morandi, Piero Parchi, Gian Luca Pirazzoli, Luisa Sambati, Caterina Tonon, Maria Giulia Bacalini

**Author notes:** Corresponding author: Francesco Ravaioli, IRCCS Istituto delle Scienze Neurologiche di Bologna, via Altura 3, 40139, Bologna, Italy. shared senior authorship (Caterina Tonon, Maria Giulia Bacalini).

## Abstract

**INTRODUCTION:** Adults with Down syndrome (DS) have a higher risk of developing Alzheimer’s disease (AD). As gut microbiota (GM) alterations have been reported in AD, we investigated their association with cognitive decline and plasma AD biomarkers in DS.

**METHODS:** Fecal and plasma samples were collected from 58 adults with DS (21-75 years) and 30 euploid controls (CTRL; 25-83 years). GM was profiled using 16S rRNA sequencing. Major Neurocognitive Disorder (NcD) was diagnosed according to DSM-5 criteria. Plasma levels of p-Tau181, NfL, and GFAP were measured using the Simoa platform.

**RESULTS:** Compared with CTRL, DS showed significant changes in *UBA1819* and *Intestinibacter* genera, previously reported to be associated with mild cognitive impairment. Furthermore, DS with NcD were characterized by a reduced abundance of *Roseburia* genus, which was also negatively associated with plasma levels of AD biomarkers.

**CONCLUSION:** Adults with DS display AD-associated changes in GM partially resembling those previously reported in euploid AD patients

## 1. Background

Down syndrome (DS) is a genetic disorder caused by complete or partial triplication of chromosome 21. DS occurs in approximately 1–2 per 1000 live births globally, with a worldwide prevalence of 1/800 [1]. Individuals with DS present an increased risk for various health issues, including congenital heart disease, developmental delay, leukemia, obesity, obstructive sleep apnea, and abnormalities in myelopoiesis and inflammatory responses [2–4]. Alterations in brain development lead to a reduction in total brain volume, particularly in the cortical, hippocampal, and cerebellar regions, and are associated with intellectual disability of variable severity [5]. Furthermore, the overexpression of the β-amyloid precursor protein (APP) gene, located on chromosome 21, increases the risk of developing Alzheimer’s disease (AD)–related dementia in adults with DS [6]. By age 40, pathological features of AD are already detectable in cerebrospinal fluid (CSF) and plasma, as demonstrated by several studies reporting elevated levels of neurofilament light protein (NfL), phosphorylated Tau (pTau181, pTau217), glial fibrillary acidic protein (GFAP), and a decline in the ratio of proteins Aβ-42/Aβ-40 [7–10]. However, the age of onset and progression of dementia in DS are highly variable, suggesting the existence of factors that can modulate the AD pathological cascade, as observed in the general population.

The gut microbiota (GM) represents the most densely populated bacterial community in the human body and plays an essential role in maintaining overall health. Intestinal bacteria and their metabolites support key functions including preserving gut barrier integrity, modulating the immune system, and regulating essential metabolic processes [11]. Moreover, the gut microbiome can influence brain function and homeostasis through the bidirectional communication network known as the gut-brain axis. This axis transmits information via neural, immune, and endocrine pathways, thereby impacting mood, cognitive functions, including memory, cognition and social behavior, and its alterations can contribute to the onset and progression of neurological and psychiatric disorders [12–16]. Several studies have suggested a link between GM alterations, AD and its preclinical conditions, reporting changes in microbial composition along the AD continuum [17–20]. Altered taxa composition may contribute to AD pathogenesis through several mechanistic pathways. Dysbiosis has been linked to elevated production of lipopolysaccharides (LPS) and microbial amyloids, which can compromise intestinal epithelial integrity and disrupt the blood–brain barrier (BBB). These alterations facilitate systemic and neuroinflammatory responses, promote oxidative stress, sustain amyloid-beta (Aβ) aggregation, induce central insulin resistance, and ultimately lead to neuronal apoptosis, all of which are characteristic features of AD neuropathology [21–23].

Studies investigating the GM in individuals with DS are limited, but available results suggest an imbalance in both microbial composition and metabolite profile compared to euploid subjects. These differences were evident even during childhood and have been associated with cognitive and behavioral disturbances [24–26]. Interestingly, a recent metagenomic study investigated fecal microbiota in a cohort of 20 DS cognitively stable or with mild cognitive impairment (MCI) and reported both overall and taxa-specific changes according to cognitive status [27].

The present study seeks to advance understanding of the role of GM in individuals with DS, with particular focus on its association with the development of AD. We investigated the fecal microbiota of adults with DS using 16S rRNA high-throughput sequencing and analysed its association with plasma levels of AD-related biomarkers [28,29], including phosphorylated pTau181, NfL, and GFAP.

## 2. Methods

### 2.1 Cohort

Participants with DS and healthy controls were recruited at AUSL Bologna -IRCCS Istituto delle Scienze Neurologiche of Bologna (Italy) in the framework of the study protocols “Study of aging in people with Trisomy 21: longitudinal evaluation of molecular markers and clinical-functional, neuropsychological and cognitive aspects (AgingInT21)” and “Digital biomarkers in Parkinson and Alzheimer diseases and in subjects with Down Syndrome (DARE-T21)”. The protocols were approved by the Local Ethics Committee of the local health service of Bologna, Italy (1070-2021-SUPER-AUSLBO and 378-2024-OSS-AULSBO) and informed written consent was obtained from the participants and from their relatives or legally authorized representatives. Geriatric and neurological evaluation was performed by expert clinicians according to a standardized protocol, as previously described [30]. Major Neurocognitive Disorder (NcD) was diagnosed according to DSM-5 TR criteria [31].

### 2.2 Sample collection and DNA extraction

Fecal samples were collected in OMNIgene®•GUT tubes to stabilize microbial DNA (DNA Genotek, Canada). Samples were stored for a maximum of 7 days at room temperature, vortexed 60 sec, transferred to cryotubes and stored at -80°C. Microbial DNA was extracted from faeces using a modified version of the protocol described by Ghosh et al [32]. Briefly, 200 ul of defrosted faeces were lysed using the ASL buffer (QIAGEN, Hilden, Germany) and bead beaten 2 times at 1800 rpm for 3 min, with a 1-minute pause between the 2 cycles, in Power Bead Tubes (Ceramic 1.4 mm) using the PowerLyzer 24 Homogenizer (QIAGEN, Hilden, Germany). After proteinase K treatment, DNA purification was performed using the QIAamp Fast DNA Stool Mini Kit (QIAGEN, Hilden, Germany). DNA was quantified using Qubit™ dsDNA Broad Range (BR) Assay Kit (Qiagen, Hilden, Germany) and stored at -20°C before analysis.

### 2.3 Analysis of plasma AD biomarkers

Venous blood was collected at fasting using sodium ethylenediaminetetraacetic acid (EDTA) tubes. Blood samples were centrifuged at 2000×g at room temperature for 10 min within 2 hours from collection. Plasma supernatant was collected, divided into aliquots, and frozen at−80 °C until use. Plasma p-Tau181, NfL and GFAP were measured with SiMOA p-tau181 advantage V2 and SiMOA Neurology 2-Plex B (GFAP, NF-l) Assay Kit, respectively. Analyses were performed on a SiMOA SR-X analyzer platform (Quanterix, Billerica, MA, USA). Biomarker levels were transformed using a base-2 logarithm.

### 2.4 Library Construction and Sequencing

Library preparation was conducted following Illumina 16S Library Preparation Workflow. Specifically, the 16S rRNA gene hypervariable V3-V4 region was amplified with primers 341F and 805R. The PCR products were purified using MagSi Prep Plus magnetic beads (Magtivio, NL) and amplified for 8 cycles using Nextera XT Indexed Primers. The final amplicons were purified, quantified using Qubit™ dsDNA BR Assay Kit (Qiagen, Hilden, Germany) and normalized at equimolar concentration (4nM). Pooled libraries were denatured with 0.2 N NaOH, diluted and combined with 10% PhiX Sequencing control V3. The final pool (6 pM) was sequenced with MiSeq v3 reagents (Illumina, San Diego, CA) on Illumina MiSeq platform using paired-end 2 x 300-bp reads.

### 2.5 Bioinformatic processing

Pre-processing of raw paired-end reads was performed according to Dada2 Big Data pipeline [33] in R (v. 4.2.2). Briefly, forward and reverse reads were checked for quality and trimmed to remove Illumina adapters and low-quality sequences. Filtered reads were deduplicated, merged and clustered into an amplicon sequence variant (ASV) table. Finally, after removal of chimeric ASVs, taxonomy was assigned against SILVA non-redundant small subunit ribosomal RNA database (v138.1) [34], with default bootstrap confidence value (50%). Samples with low sequencing depth (<10000 reads) were excluded from the analysis (88 samples remaining). Taxa that were assigned to neither Bacteria nor Archaea and taxa present in less than 10% of the total sample size were removed from the ASVs count table.

### 2.6 Statistical analyses

All statistical analyses were performed with R (v. 4.2.2). Age and sex were included in all parametric statistical models as confounding variables. For analyses performed within the DS group, Body Mass Index (BMI) was included as a covariate (correction was not applied to euploid controls due to unavailability of BMI data). Statistically significant threshold was considered at nominal p-values<0.05; in addition, false discovery rate (FDR) according to the Benjamini-Hochberg (BH) method was calculated Alpha and beta diversity at ASVs level were calculated using the *phyloseq* package (v 1.42.0) [35]. Alpha diversity was calculated according to the number of observed species, Chao, Shannon and inverted Simpsons indexes. Wilcoxon rank-sum test was used to compare alpha diversity measurements between DS and CTRL groups, as alpha diversity data was not normal according to the Shapiro-Wilk test. The association between alpha diversity metrics and levels of plasma markers p-Tau181, NfL and GFAP in DS subjects was evaluated by fitting a linear regression model. Beta diversity was assessed via Principal Component Analysis (PCoA) based on Bray-Curtis dissimilarity, weighted and unweighted Unifrac metrics. PERMANOVA function in *vegan* R package (v 2.6-6.1) was used to compare beta-diversity metrics between DS and CTRL participants. Alterations in microbial abundances were evaluated at phylum, family and genus level. Firstly, dysbiosis was evaluated by calculating Firmicutes/Bacteroidetes ratio and compared between DS and CTRL participants using ANOVA test. Secondly, DESeq2 pipeline [36] using Wald test was used to perform differential microbial abundance analyses between DS and CTRL, as well as between DS with and without NcD. An alternative analysis of differential abundance was performed using the Aldex2 method [37]. The association between microbial abundance and AD plasma biomarkers was also assessed using the DESeq2 pipeline. Finally, predictive functional profiling was performed using PICRUSt2 (v. 2.6.2) [38]. Predicted abundances were normalized with Centered Log-Ratio (CLR) transformation using the compositions package (v. 2.0-6). Differential pathway abundance analysis in DS versus CTR and in DS with NcD versus DS without, as well as the association of pathway abundance with AD biomarkers, were conducted in R using limma package (v. 3.54.2).

## 3. Results

### 3.1 Demographic characteristics of participants

After quality filtering, metagenomic data were available for 58 adults with Down Syndrome (DS; 20 females, mean age 42 years) and 30 euploid controls (CTRL; 24 females, mean age 58 years). Nine DS participants had a diagnosis of Major Neurocognitive Disorder (NcD), while 5 had chronic gastrointestinal diseases. Mean value of Body Mass Index (BMI) in DS was 27.7 (13 with BMI>30, obese). BMI values were not available for CTRLs. Demographic and clinical characteristics of the cohort are summarized in **Table 1**.

**Table 1.** Demographic characteristics of participants. Abbreviations: SD, corresponds to standard deviation; n, number; DS, Down Syndrome participants; CTRL, euploid subjects; *p*, p-value. *, calculated by t-test; †, calculated by Fisher test.

|  | CTRL group (n= 30) | DS group (n= 58) | *p* |
| --- | --- | --- | --- |
| *Demographic* |  |  |  |
| Age (mean years, range) | 58 (25-83) | 42 (21-75) | <0.001* |
| Females (n/total) | 24/30 | 20/58 | <0.001† |
| BMI (mean ±SD) | / | 27.7±5.9 | / |
| *Dementia clinical diagnosis* |  |  |  |
| Major Neurocognitive Disorder (n) | / | 9 | / |
| *Comorbidities (n)* |  |  |  |
| Gastrointestinal disorders | / | 5 | / |
| Type 2 Diabetes | / | 0 | / |

### 3.2 Plasma AD biomarkers

abundance with AD biomarkers, were conducted in R using *limma* package (v. 3.54.2).

Plasma levels of AD biomarkers (pTau181, NfL and GFAP) were available for 51 DS (19 females; mean age 40 years; 9 with a diagnosis of NcD) and 12 CTRL (8 females; mean age 40 years). As expected [29], biomarker levels were higher in DS compared to CTRL (ANOVA correcting for age and sex; pTau181: p-value=0.018; NfL: p-value<0.001; GFAP: p-value=0.001; **Figure 1A**) and showed a relevant increase after 40 years in the DS group **(Figure 1B).** Furthermore, pTau181 and NfL were significantly higher in DS with NcD than in DS without NcD (ANOVA correcting for age and sex; pTau181: p-value=0.025; NfL: p-value=0.017; **Figure 1C)**.

**Figure 1.**
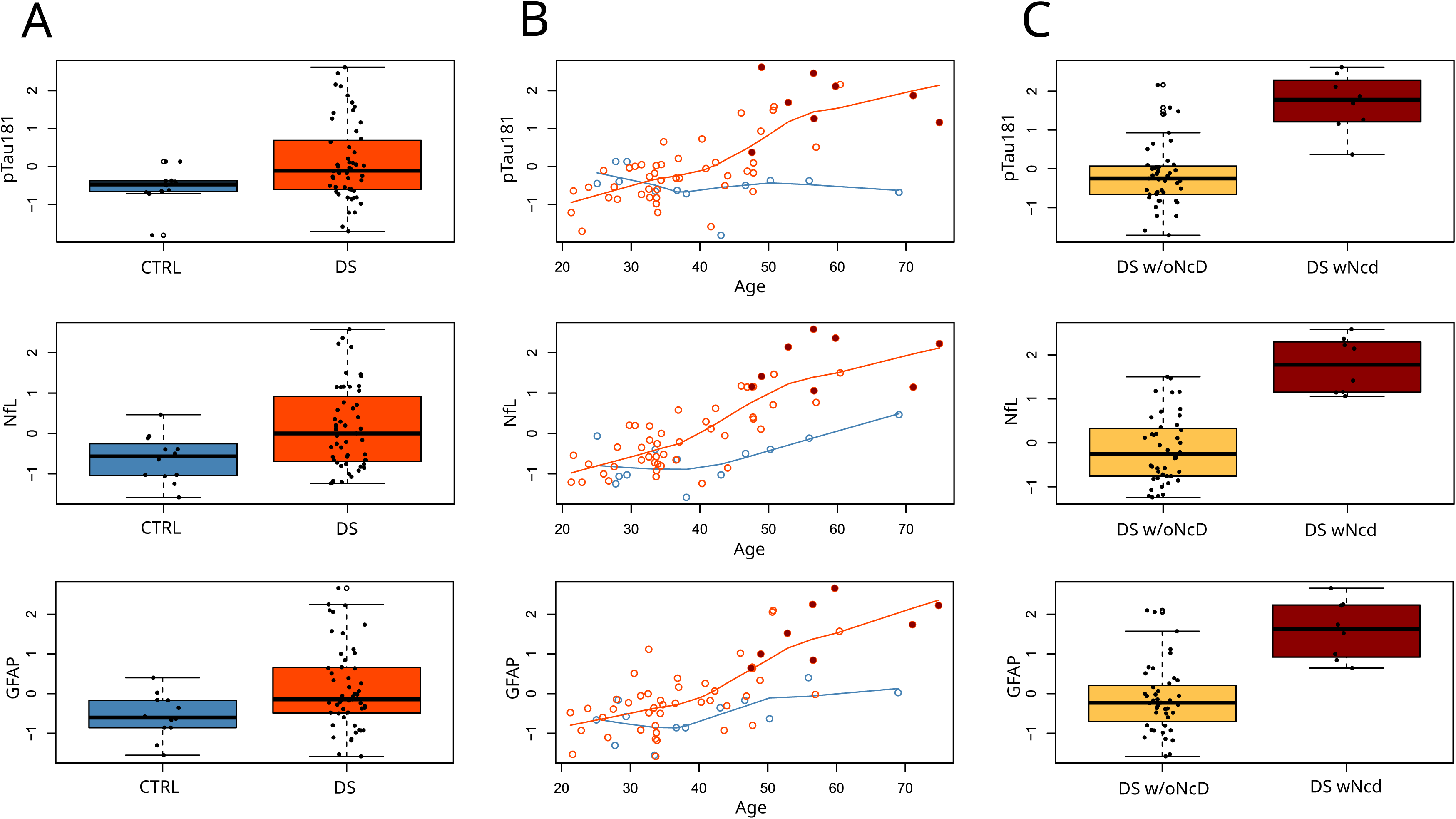
Plasma levels of p-Tau181, NfL and GFAP. **A**. Boxplots of pTau181, NfL and GFAP levels in DS (red) and CTRL (blue) groups. Sample size n=63. **B**. Scatterplots of pTau181, NfL and GFAP levels with respect to the age of DS (red) and CTRL (blue); DS with NcD are colored in dark red. Sample size n=63. **C**. Boxplots of pTau181, NfL and GFAP levels in DS without NcD (w/oNcD, yellow) and DS with NcD (wNcD, dark red). Plasma biomarker levels are reported as z-scores of base-2 logarithm-transformed values. Sample size n=51.

### 3.3 Analysis of gut microbiota in DS and CTRL groups

The 16S rRNA amplicon sequencing yielded a total of 2284703 reads (ranging from 11177 to 57267 per sample) passing quality filters. After prevalence filtering, the dataset comprised 275 ASVs that were assigned to 6 phyla, 21 families and 62 genera.

As a first step, we evaluated differences between CTRL and DS groups in terms of overall microbiota diversity (alpha and beta-diversity) and microbial abundance at phylum, family and genus levels.

Alpha diversity, which is a within-sample measure of microbiota diversity, did not show significant differences between DS and CTRL groups (Wilcoxon test p-value>0.05; **Figure 2A** and **Supplementary File 1**). The analysis of beta diversity, which measures microbial community dissimilarities among groups, showed significant differences between DS and CTRL groups with Bray-Curtis distance (p-value=0.021), but not with Weighted and Unweighted Unifrac metrics (**Figure 2B** and **Supplementary File 1**).

**Figure 2.**
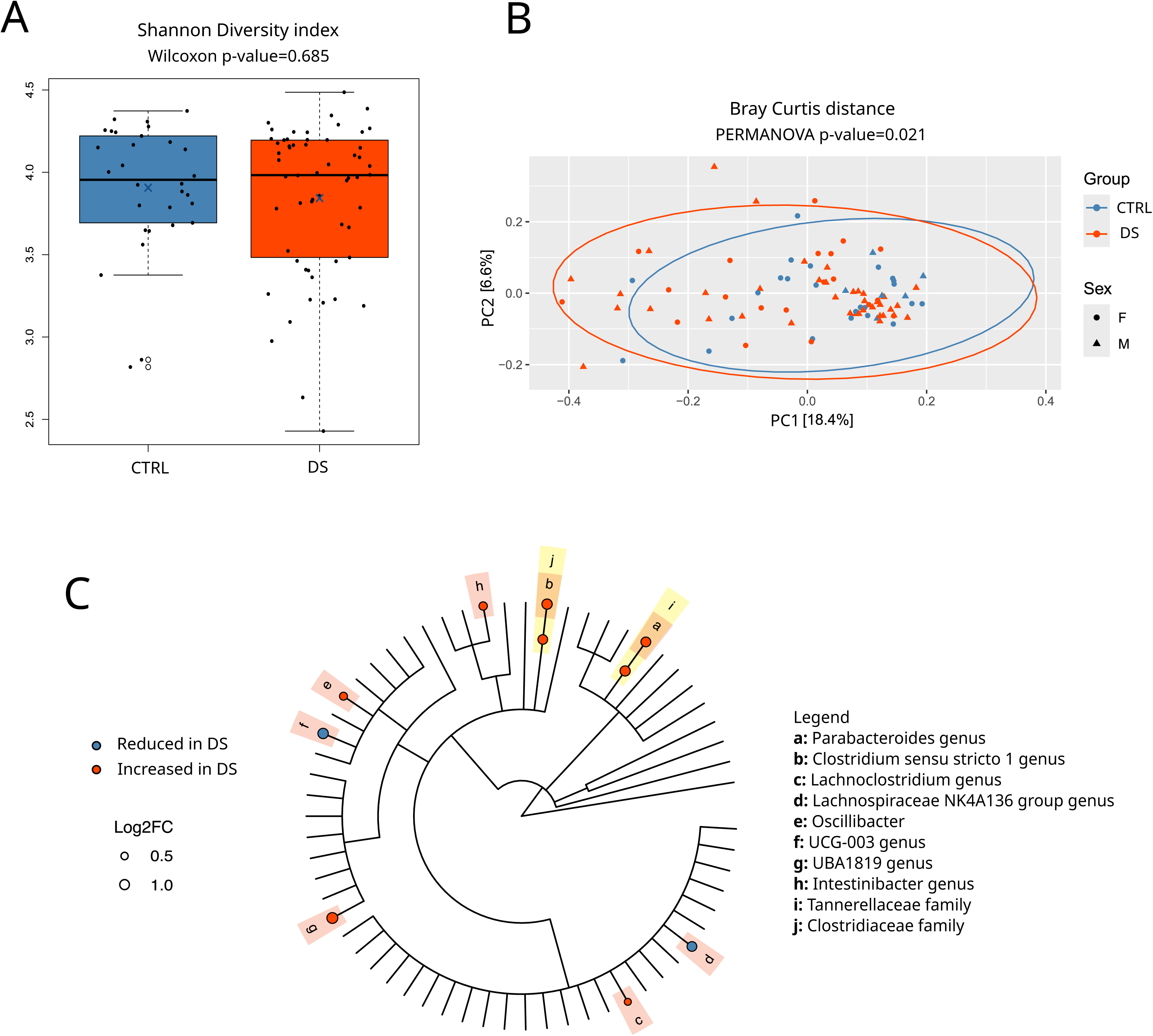
Microbial diversity and taxa abundance differences between the DS and CTRL groups. **A**. Boxplot showing alpha-diversity values calculated according to Shannon index in DS and CTRL groups. Group means are indicated with a blue X symbol; group medians are indicated with a continuous black line. Sample size n=88. **B**. Beta diversity plot showing separation of DS (red) and CTRL (blue) groups based on Bray-Curtis dissimilarity metric. Sample size n=88. **C**. Cladogram showing different abundant taxa levels in the DS group as identified by Deseq2 analysis. Microbial families showing significant changes are highlighted in yellow, while genera are highlighted in orange. Blue and red dots indicate taxa significantly reduced and increased in the DS group, respectively. The dimension of each dot is proportional to the Log2 Fold Change (Log2FC). Sample size n=88.

We used DESeq2 to analyze differential microbial abundance in DS compared to CTRL participants at different taxonomic levels (phylum, family and genus) (**Supplementary File 1**).

The analysis of bacterial phyla abundance revealed no statistically significant differences between groups. Furthermore, the ratio of Firmicutes to Bacteroidetes (F/B ratio) was not altered in DS compared to CTRL (**Figure 2C** and **Supplementary File 1**). At the family level, we found that the abundance of *Tennerellaceae* and *Clostridiaceae* was significantly higher in DS than in CTRL (nominal p-value: 0.012, 0.012, respectively; **Figure 2C** and **Supplementary File 1**). Finally, we identified 8 genera that showed significant abundance alterations in DS compared to CTRL **Figure 2C** and **Supplementary File 1**): *UCG-003* (nominal p-value= 0.039) and *Lachnospiraceae NK4A136 group* (nominal p-value= 0.002) were reduced in DS, while *Lachnoclostridium* (nominal p-value= 0.044)*, Oscillibacter* (nominal p-value= 0.021)*, Intestinibacter* (nominal p-value= 0.020)*, Parabacteroides* (nominal p-value= 0.025)*, Clostridium sensu stricto 1* (nominal p-value= 0.004) and *UBA1819* (nominal p-value<0.001) genera were increased in DS. For *UBA1819* statistical significance survived also after false discovery rate (FDR) correction (q-value=0.028) (**Figure 2C** and **Supplementary File 1**).

To confirm the above-described results, we used an alternative analytical approach based on *Aldex2* package. This analysis confirmed the alterations between DS and CTRL for *Parabacteroides, Intestinibacter* and *Lachnospiraceae NK4A136 group* genera (nominal p-value= 0,029; 0,020 and 0,043, respectively). Moreover *Lachnospiraceae UCG-010, Clostridium sensu stricto 1 and UBA1819* were marginally significant (nominal p-value= 0,056; 0,059 and 0,059, respectively) (**Supplementary File 2**).

Finally, functional abundance analysis performed using PICRUSt2 tool identified 32 significant pathways in the comparison between DS and CTRL groups (**Supplementary File 3**). Considering the higher-level category annotations, we found that DS showed a reduction in several significant pathways related to quinol and quinone biosynthesis (12/32), including menadione (vitamin K2) biosynthesis.

### 3.4 Gut microbiota analysis according to clinical diagnosis of Major Neurocognitive Disorder

To evaluate a possible association between GM composition and cognitive status in adults with DS, we compared DS subjects with and without a clinical diagnosis of NcD.

Analysis of alpha diversity did not reveal significant differences for any index, while beta diversity calculated with Unweighted UniFrac index differed between DS with and without NcD (**Figure 3A** and **B** and **Supplementary File 4**). In the analysis of taxa abundance, the *Firmicutes* phylum abundance was significantly reduced in DS with NcD (nominal p-value= 0.048), while no significant differences emerged in the F/B ratio (data not shown). At the family level, DS with NcD showed a reduction of *Peptostreptococcaceae* (nominal p-value= 0.023) and an increase of *Tannerellaceae* (nominal p-value=0.027) (**Figure 3C** and **Supplementary File 4**). At genus level, DS with NcD showed a decrease of *Roseburia* (nominal p-value=0.004) and *Lachnospiraceae UCG-010* (nominal p-value=0.005), and an increase of *Parabateroides* (nominal p value=0.017), *Alistipes* (nominal p-value=0.037) and *Butyricicoccus* (nominal p value=0.039) (**Figure 3C** and **Supplementary File 4**). Functional abundance analysis identified 2 significant pathways (Biosynthesis of Quinol and Quinone, and Fatty Acids and Lipid Biosynthesis classes), which were increased in DS with NcD (**Supplementary File 3**).

**Figure 3.**
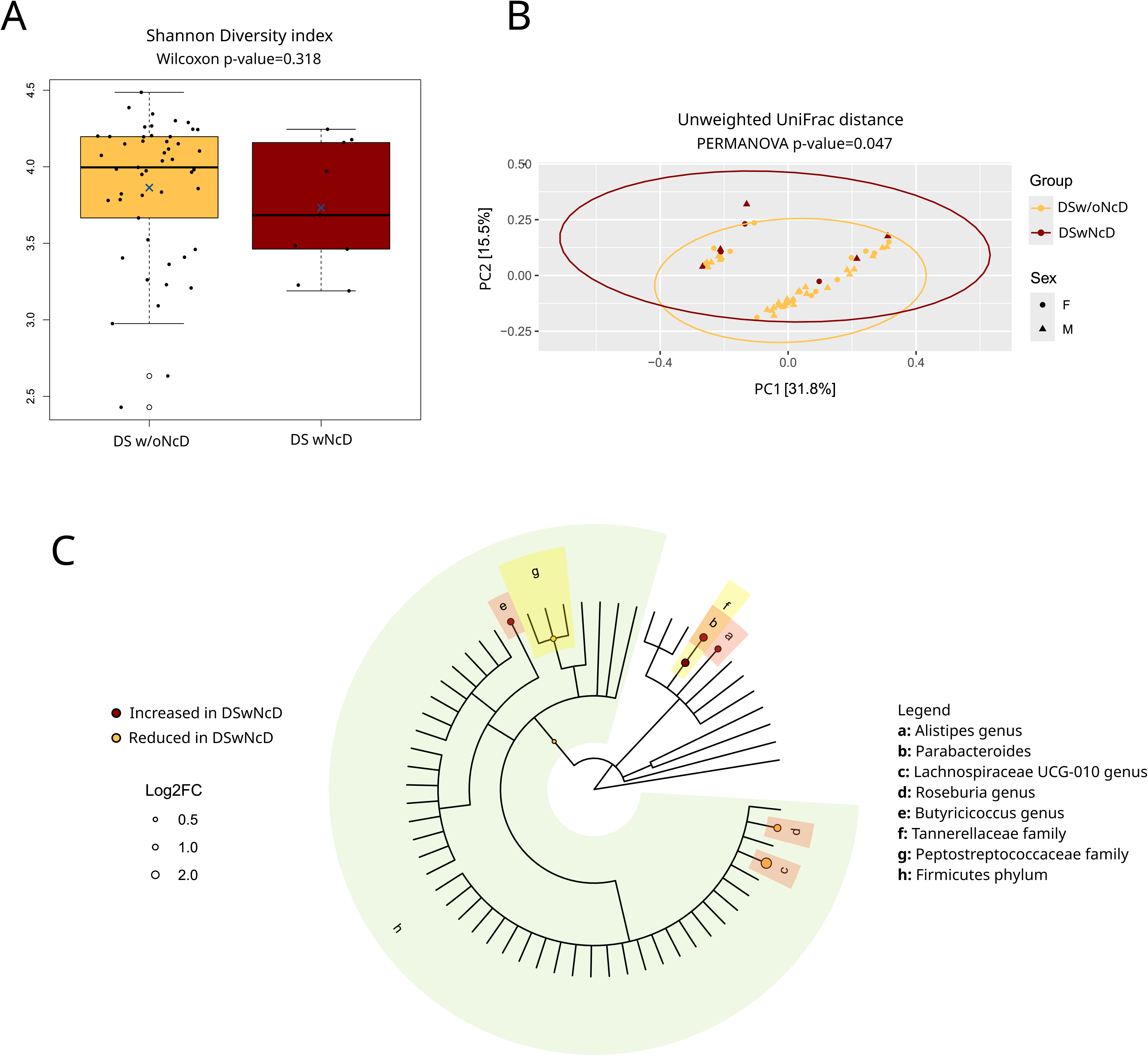
Microbial diversity and taxa abundance differences between DS with and without NcD. **A**. Boxplot showing alpha-diversity values calculated according to Shannon index in DS wNcD and w/oNcD groups. Group means are indicated with a blue X symbol; group medians are indicated with a continuous black line. Sample size n=58. **B**. Beta diversity PCoA plot showing separation of DS wNcD (dark red) and DS w/oNcD (yellow) groups based on unweighted UniFrac metric. Sample size n=58. **C**. Cladogram showing the differentially abundant taxa levels between DS wNcD and w/oNcD groups as identified by DeSeq2 analysis. Microbial phyla showing significant changes are highlighted in green, families in yellow, while genera are highlighted in orange. Taxa significantly increased in the DS wNcD group are indicated with dark red dots, while those reduced in DS wNcD in yellow. The dimension of each dot is proportional to the Log2 Fold Change (Log2FC). Sample size n=58.

### 3.5 Gut microbiota analysis according to plasma AD biomarkers

Finally, we analyzed the association between GM composition and plasma AD biomarkers in DS participants. No significant associations were found between alpha diversity metrics and plasma levels of p-Tau181, NfL or GFAP. Conversely, we detected significant associations for specific genera (**Figure 4A** and **Supplementary File 5**). In particular, we found a significant association with p-Tau181 for the 3 genera *Roseburia* (nominal p-value= 0.033)*, Lachnospira* (nominal p-value= 0.020) and *UBA1819* (nominal p-value= 0.009); a significant association with NfL for the 2 genera *Roseburia* (nominal p-value= 0.035) and *Christensenellaceae R-7 group* (nominal p-value= 0.013); and a significant association with GFAP for the 2 genera *[Eubacterium] siraeum group* (nominal p-value= 0.005) and *Lachnospiraceae UCG-001* (p-value= 0.014). Notably, *Roseburia* showed a significant negative association with both p-Tau181 and NfL, and it showed a marginally significant association with GFAP (nominal p-value= 0.055) (**Figure 4B** and **Supplementary File 5**).

**Figure 4.**
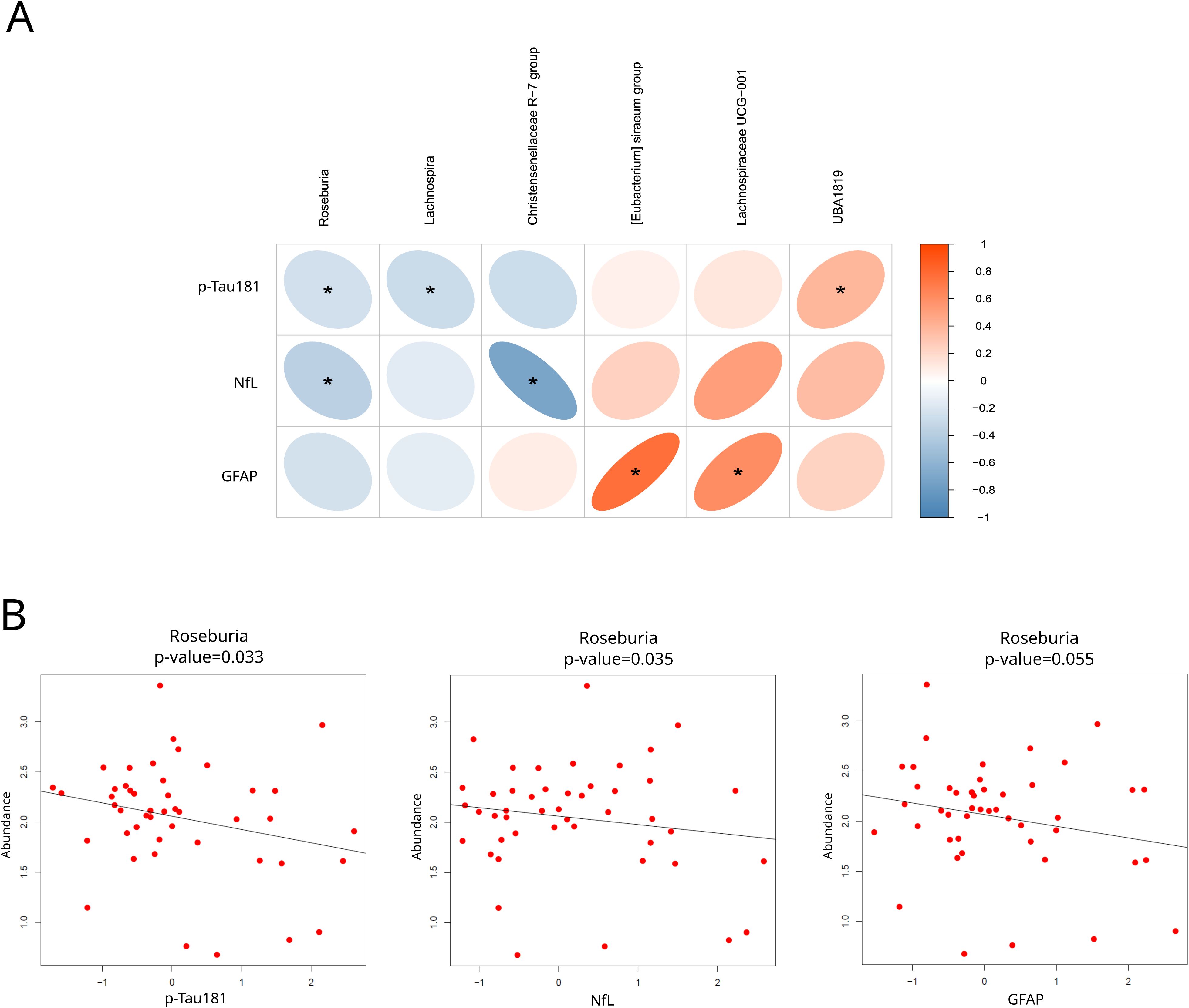
Association analysis of genera abundance and AD plasma biomarkers. **A**. Correlation matrix between plasma AD biomarkers (p-Tau181, NfL and GFAP) and genera abundance in DS participants. The matrix reports only the genera that show a significant association with at least one biomarker. Colors indicate Pearson correlation coefficient values. The elliptical shape indicates the strength of correlation between microbial genus and plasma biomarker. In the correlation visualization, bolder colors indicate higher correlations. If the shape of an ellipse bends towards the right, it indicates positive correlation, whereas negative correlation if its shape bending towards the left shows a negative correlation. Sample size n=47. **B**. Scatterplots reporting an example of the results obtained from the association with AD plasma biomarkers levels. *Roseburia* abundance is represented in association with p-Tau181, NfL and GFAP plasma levels. Sample size n=46.

Functional abundance analysis identified no pathway significantly associated with p-Tau181, whereas 55 and 11 pathways emerged as significantly associated with NfL and GFAP, respectively (**Supplementary File 3**). In particular, we found several pathways annotated within the Proteinogenic Amino Acid Biosynthesis (4/55), Enzyme Cofactor Biosynthesis (8/55) and Alcohol Degradation (4/55) classes that were negatively associated with NfL. In addition, we identified several pathways annotated within the Fatty Acids and Lipid Biosynthesis (4/55) classes that were positively associated with NfL. Notably, several pathways related to Fermentation to Short-Chain Fatty Acids class that were significantly reduced in the DS group were also negatively associated with NfL and GFAP plasma levels (**Supplementary File 3**).

## 4. Discussion

In this study, we analyzed the gut microbiome composition in adults with DS in comparison with euploid controls, and we further examined how microbiome variation relates to clinical diagnosis of NcD and to the levels of plasma AD biomarkers.

Our data suggest no major differences in the overall gut microbiota composition of adults with DS compared to euploid subjects, in accordance with what was previously reported by Biagi et al. in a cohort having demographic characteristics similar to ours [24]. In particular, we found that the DS group exhibited weak but significant differences in beta-diversity based on Bray–Curtis metric, whereas no variations in alpha-diversity metrics were observed. Conversely, differences in alpha diversity with respect to euploid controls were previously reported in children and young adults with DS [25,26], although with a discordant direction of change in the two available studies. These heterogeneous results can be ascribed to differences in the demographic characteristics of the cohorts evaluated in our and previous studies, limited sample sizes, and variations in experimental and analytical pipelines. Similarly, there is no overlap in the results of differential abundance analysis between DS and CTRL groups across the different studies, including ours. Despite the similarities in the demographic characteristics of the cohorts, a direct comparison with Biagi et al. is not possible, as our analytical pipeline filtered out the low prevalence genera reported there as significant (*Parasporobacterium*, *Sutterella* and *Veillonellaceae*) [24]. Functional abundance analysis in our dataset identified a reduction of pathways related to menaquinone (Vitamin K2) biosynthesis in DS. Menaquinone, mainly produced by the GM, is known to improve bone health and prevent coronary calcification, but several studies have also reported a key role in preserving cognitive functions [39] and brain health [40]. Animal studies showed that menaquinone administration can reverse cognitive changes induced by antibiotic-induced gut dysbiosis, pointing to a neuroprotective role mediated by a reduction of oxidative stress and inflammation in intestine and brain [41]. Therefore, we can hypothesize that the GM changes that we observed in our dataset may contribute to the enhanced neuroinflammation characteristic of DS [42].

As adults with DS are genetically at higher risk of developing AD at an early age, it is of interest to investigate whether the GM alterations described in euploid AD patients are already present in our entire DS cohort, which includes participants of different ages and with and without cognitive decline. Among the genera identified by differential abundance analysis, *Clostridium sensu stricto 1* and *Intestinibacter* were previously reported to be decreased in AD patients [43,44], while they showed the opposite direction of change in our DS cohort. Conversely, *UBA1819* and *Intestinibacter* were reported to be elevated in euploid subjects with a clinical diagnosis of MCI in a Greek cohort [45], consistent with our findings. These results suggest that, when considering our entire cohort, the observed GM alterations show some similarities with those reported in MCI, but not with those described in AD patients. Within this framework, it is worth noting that some studies suggest the presence of stage-specific GM changes along the AD continuum in the general population, with alterations observed in AD being only partially detected in patients with clinical MCI and vice-versa [46,47].

We then specifically evaluated the GM alterations associated with Major Neurocognitive Disorder in DS. To the best of our knowledge, only Rosas et al. recently investigated this topic, comparing 14 cognitively stable DS and 6 DS with MCI, designed on the basis of the assessment of subtle cognitive and/or functional decline [27]. Our dataset included a clinical characterization (diagnosis of major NcD by DSM-5 criteria) and a biological characterization (plasma biomarkers) of AD associated cognitive decline.

When comparing participants with DS with and without NcD, we found a slight but significant difference in beta diversity and a reduction of *Firmicutes* phylum in the NcD group, in accordance with the results reported by Rosas et al. [27]. *Firmicutes* phylum includes many SCFAs producers with protective and anti-inflammatory properties [48], and its reduction has been reported in AD [20,49]. Our differential abundance analysis revealed additional taxa previously reported to be altered in euploid AD patients. Among them, there are the *Peptostreptococcaceae* family, reduced in the NcD DS group as in AD [50], and the *Alistipes* and *Roseburia* genera, respectively increased and decreased in the NcD group, which have been consistently reported as altered also in AD and associated with cognitive decline [18,51–53].

The analysis of the association between taxa abundance and plasma AD biomarkers in participants with DS further expanded the results described above. Remarkably, we found that the abundance of *Roseburia* genus showed a negative association with all the biomarkers, confirming similar analyses performed on CSF AD biomarkers (amyloid-β1-42 and p-tau) in euploid patients [18]. The latter study also identified *Christensenellaceae R7 group* genus, which was negatively associated with NfL in our dataset and was also reported to be reduced in DS with MCI by Rosas et al [54]. In addition, *UBA1819* genus, which we found significantly increased in DS compared to euploid controls, was positively associated with p-Tau181.

In summary, our results extend previous findings on GM alterations and cognitive decline in DS and highlight for the first time their association with plasma AD biomarkers. These results require further validation in larger cohort studies and should be complemented by additional characterizations, including fecal metabolomic profiling, to better elucidate the functional implications of the microbial changes that we observed in the pathogenesis and progression of AD in adults with DS. This line of investigation holds substantial promise for future research and may identify novel targets for early intervention against AD in adults with DS.

## Abbreviations

CTRL: Control euploid subjects
DS: Down syndrome
AD: 2 Alzheimer’s disease
NcD: Major Neurocognitive Disorder
CNS: Central Nervous System
GM: Gut microbiota
CTRL: Control euploid subjects
DS: Down syndrome
AD: 3 Alzheimer’s disease
NcD: Major Neurocognitive Disorder
CNS: Central Nervous System
GM: Gut microbiota
BBB: Blood–brain barrier
LPS: Lipopolysaccharides
SCFAs: Short-chain fatty acid
MCI: Mild Cognitive Impairment

## Acknowledgements

The authors thank all the people with Down syndrome who participated in the study and their families. The authors thank Marica Lanzoni, Claudia Boninsegna, Elisabetta Venieri and Silvia De Luca (IRCCS Istituto delle Scienze Neurologiche di Bologna, Bologna, Italy) for their assistance in planning and organizing the study operational procedures.

## Consent Statement

All participants gave their written informed consent for the use of their biological material and clinical data for research purposes. Ethical approval was granted by the Local Ethics Committee of the local health service of Bologna.

